# Simplified In Vitro Generation of Human Gastruloids for Modelling Early Development

**DOI:** 10.1101/2025.10.15.682610

**Authors:** Takuya Azami, E. Elizabeth Patton, Jennifer Nichols

## Abstract

The purpose of this study was to optimize the efficiency and cost of human gastruloid formation by testing and adjusting individual parameters using as examples two distinct human pluripotent stem cell lines, both available from the UK Stem Cell Bank. For the first step, commercially sourced culture medium was replaced with a home-made defined recipe, known as ‘N2B27’, into which specific reagents can be titrated. By reducing the concentration of Activin A to 15% of the original protocol, efficient elongation of aggregated embryoid bodies was achieved. Also, titrating initial cell density and delaying the brief culture in GSK3 inhibitor until the onset of cell aggregation in individual wells was advantageous. Efficiency of formation of early gastruloids exhibiting the expected regionalization of the three embryonic germ layers was further enhanced by addition of TGF*β*-inhibitor. The optimization steps presented here thus provide a simplified, robust and relatively economical protocol for consistent generation of elongated gastruloids from human pluripotent stem cells. This streamlined method improves accessibility and reproducibility, also providing a standardized platform to investigate fundamental principles of early human development.

**Summary Statement:** We present an optimized protocol for human gastruloid production that should enhance efficiency, reproducibility and affordability for future *in vitro* studies into early human post-implantation development.

## Introduction

Human gastrulation begins with the differentiation of primitive streak (PS) cells around embryonic day 14–15, Carnegie stages (CS) 6–7, and concludes by approximately day 21 (CS9), culminating in the formation of the three germ layers: ectoderm, mesoderm, and endoderm (Hertig et al., 1956). This developmental phase is also characterized by the spatial organization of distinct cell types along the embryonic body axes. Nevertheless, our understanding of gastrulation in human remains relatively limited owing to the practical and ethical inaccessibility of early postimplantation human embryos (Rossant, 2024). Gastruloids, three-dimensional models for early mammalian postimplantation development and gastrulation, first developed using mouse embryonic stem cells (mESCs) (van den Brink et al., 2014), enable study of axial elongation and germ layer differentiation. Gastruloids made from human pluripotent stem cells (hPSCs) provide an exciting system with which to investigate human development post-implantation, but required modification to the original mouse protocols and optimization for individual hPSC lines (Moris et al., 2020; Olmsted and Paluh, 2021).

In the case of mouse gastruloids, naïve pluripotent mESCs are aggregated and differentiated to a postimplantation epiblast-like state for the first 48 hours of the protocol (Beccari et al., 2018; Turner et al., 2017). Gastrulation is then stimulated using the GSK3*β* inhibitor, CHIR99021 (CHIR), which in this context acts as a WNT agonist. In contrast, conventional hPSCs are generally maintained in a ‘primed’ pluripotent state, representative of the postimplantation epiblast, rendering them rapidly responsive to gastrulation-inducing signals. The original protocols pioneering formation of human gastruloids depended upon pre-CHIR treatment (Hamazaki et al., 2024; Moris et al., 2020; Neupane et al., 2025; Olmsted and Paluh, 2021); however, how CHIR pretreatment impacts gastruloid generation and reproducibility has not been determined. In this study, we set out to optimize and simplify the conditions for human gastruloid formation. We modified the protocol to include ‘in house’ formulated medium in which we tested multiple variables using two distinct hPSC lines with the aim of improving germ layer specification and consistency. We demonstrate that cell density during CHIR stimulation critically influences gastruloid induction, particularly the balance between FOXA2⁺ endoderm and TBXT⁺ mesoderm populations. Low-density cultures favour FOXA2⁺ endoderm differentiation, whereas high-density conditions suppress endodermal specification. Based on these findings, we have established a human gastruloid protocol independent of the pre-CHIR treatment step using defined culture conditions. This improved method enables highly reproducible generation of robust elongated gastruloids with the added advantage of reducing cost by replacing commercial medium with N2B27 formulated in-house as well as significantly reduced concentrations of Activin A.

## Results and Discussion

### Low Activin A enhances human gastruloid generation and consistency

We first attempted to generate human gastruloids using a defined culture medium: N2B27, instead of the commercially produced Nutristem or Essential 8, supplemented with Activin A (20 ng/ml), FGF2 (10 ng/ml), and XAV939 (WNT inhibitor, 2 µM) (hereafter referred to as ‘A20FX’) (Kinoshita et al., 2020; Rostovskaya et al., 2019). Our formulation enables modulation of signalling pathways by adding or withdrawing individual components. We used capacitated primed (p) HNES1 (Guo et al., 2016; Rostovskaya et al., 2019) and H9 (Thomson et al., 1998) hPSCs, both cultured in A20FX for at least three passages, for generating gastruloids. As cell density during the pre-CHIR treatment prior to aggregation has been shown to be advantageous (Moris et al., 2020), we plated cells at various densities in 12-well plates according to our initial culture regime (Fig. 1A). Under these conditions, A20FX-cultured hPSCs produced disorganised and degenerating gastruloids at all tested plating densities (Fig. 1B, C). We titrated the Activin A concentration to 3 ng/ml (A3FX), since capacitated hPSCs can be maintained efficiently in low Activin A containing medium (Rostovskaya et al., 2019) and low Activin A concentration enables maintenance of pluripotency of hPSCs or induced (i)PSCs (Tomizawa et al., 2011; Xiao et al., 2006). Elongated gastruloids were formed from both HNES1 (Guo et al., 2016) and H9 (Thomson et al., 1998) cells in A3FX culture; however, the efficiency was dependent on plating density, with the highest efficiency observed at 6x10⁴ cells per 12-well plate (Fig. 1B, C, and D). These experiments confirmed the importance of optimising cell density prior to pre-CHIR treatment for consistent formation of elongated human gastruloids.

**Figure 1.**
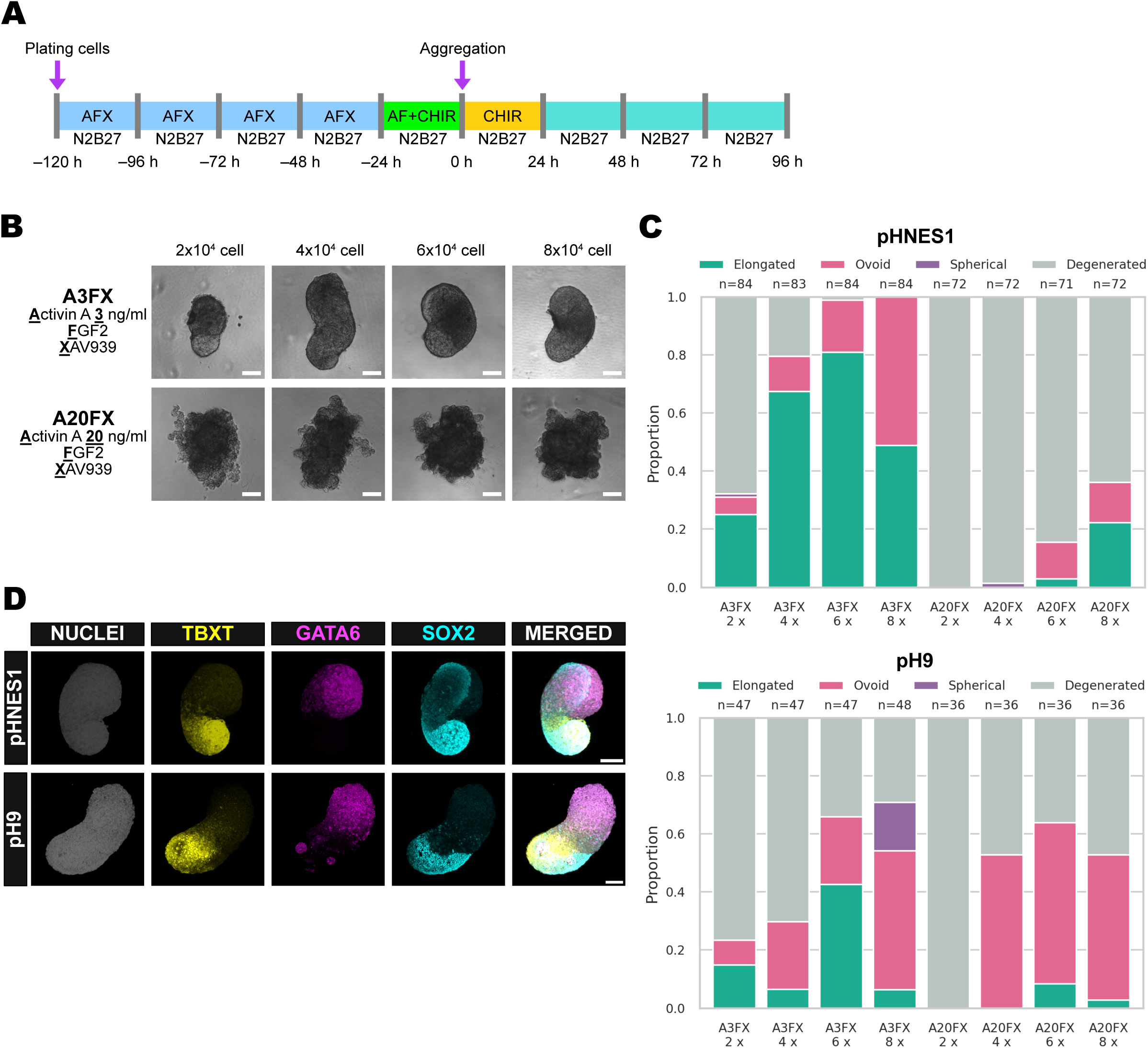
Activin A concentration and cell culture density determine the efficiency of elongated gastruloids formation. (A) Schematic of protocol for making human gastruloids, adapted from (Moris et al., 2020). (B) Representative bright field image for gastruloids generated from A3FX or A20FX cultured pHNES1 cells plated at 2x, 4x, 6x, and 8x10^4^ cell/12-well plate. Scale bar = 100 µm. (C) Proportion of elongated, ovoid, spherical, and degenerated morphology from pHNES1 and pH9 hPSCs. Plot represents a pool of three independent experiments. The number of gastruloids (n) for each condition is indicated above the corresponding plot. (D) Immunofluorescence for TBXT, GATA6, and SOX2 in gastruloids generated by indicated conditions. Scale bar = 100 µm.

### The balance between proportions of TBXT, SOX2, and FOXA2 positive cells is critical for human gastruloids

Since cell density prior to pre-CHIR treatment appeared critical to form elongated human gastruloids, we stained cells at various densities after pre-CHIR treatment to estimate the relationship between cell density and gastruloid formation (Fig. 2A, B). Expression of the endoderm marker, FOXA2, was preferentially induced in low density culture (2 or 4 x 10^4^ cells); TBXT and SOX2 expressing cells were distributed randomly, whereas at high-density, FOXA2 expression was not induced and TBXT positive cells were located at the edge of colonies, with SOX2 positive cells remaining in the middle (Fig. 2C; Fig. S1A). This expression pattern is similar to 2D micropattern gastruloids, localising SOX2 at the centre and TBXT at the periphery (Etoc et al., 2016; Martyn et al., 2019a; Martyn et al., 2019b; Yoney et al., 2018). Although both micropatterns and pre-CHIR treatment induce gastrulation, the micropattern protocol includes BMP4 to activate WNT signalling. In high-density hPSC colonies, cellular responses to TGF-*β* ligands were spatially restricted to the colony edge. This is because dense epithelialisation leads to the re-localisation of TGF-*β* receptors to the basolateral surface, where they are shielded from apical ligands by tight junctions (Etoc et al., 2016). Both FOXA2 and TBXT expression were suppressed by high density culture in A3FX (Fig. 2C, D; Fig. S1A); TBXT and LEF1 were reduced as SOX2 was reciprocally increased with higher cell density (Fig. 2E, F).

**Figure 2.**
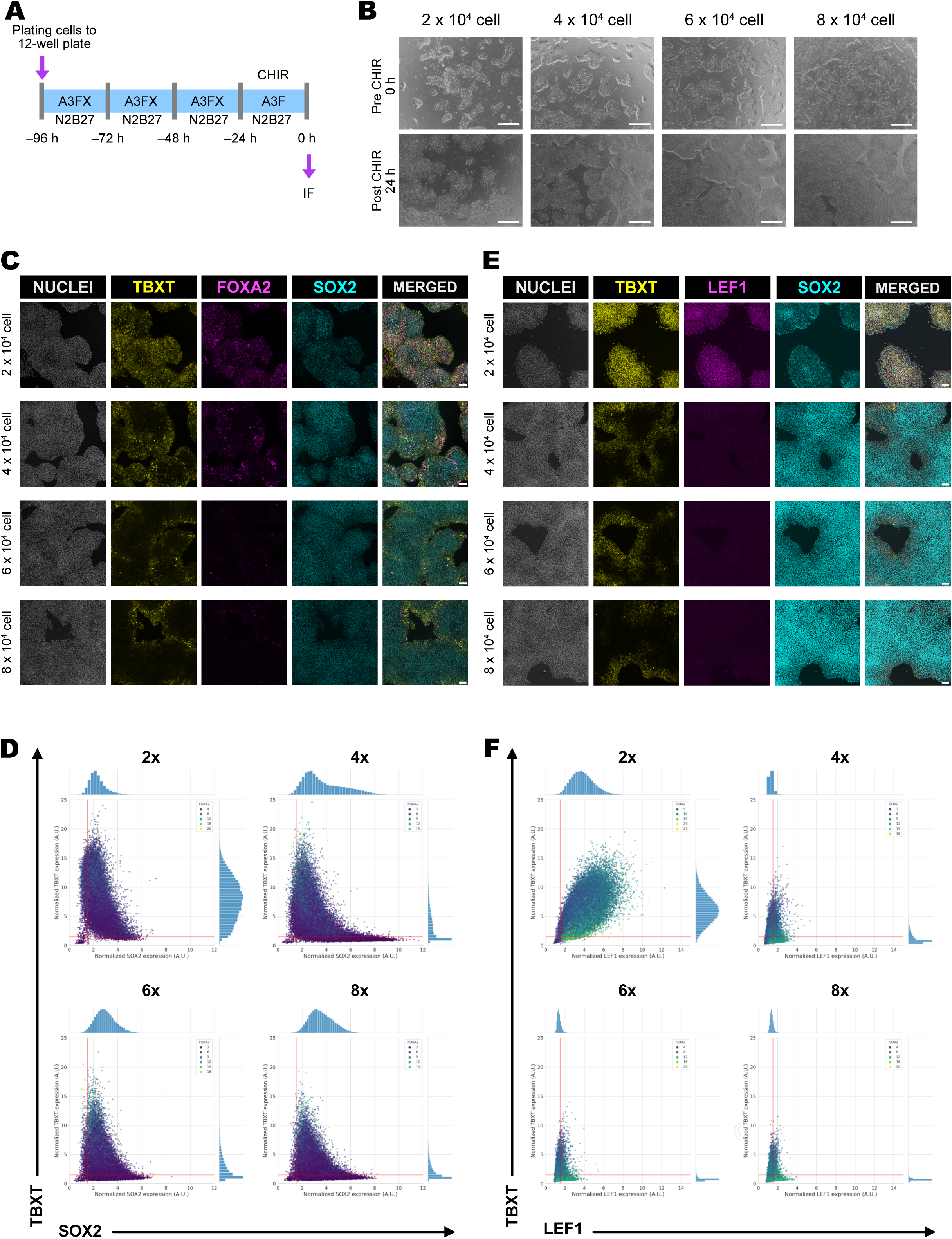
Culture density influence on the balance on TBXT, SOX2, and FOXA2 expression. (A) Schematic for the culture and pre-CHIR treatment of hPSCs. (B) Representative bright field images of pHNES1 PSCs before and after pre-CHIR. Scale bar = 100 µm. (C) Immunofluorescence for TBXT, FOXA2, and SOX2 in pre-CHIR treated pHNES1 PSCs. Scale bar = 100 µm. (D) Scatter plot for TBXT and SOX2 expression analysed from the staining in (C). Mean signal intensity was quantified from 20 fields. Heatmap showing FOXA2 expression levels. The lines on the x and y axes represent the positivity threshold, which was set based on the background fluorescence from a secondary antibody-only control. (E) Immunofluorescence for TBXT, LEF1, and SOX2 in pre-CHIR treated pHNES1 PSCs. Scale bar = 100 µm. (F) Scatter plot for TBXT and LEF1 expression analysed from the staining in (E). Mean signal intensity was quantified from 20 field. Heatmap showing SOX2 expression levels. The lines on the x and y axes represent the positivity threshold, which was set based on the background fluorescence from a secondary antibody-only control.

FOXA2 was barely visible in A20FX at high density, and TBXT was reduced as density increased (Fig. S1B). These data indicate that high cell density in culture conditions prior to CHIR treatment maintains cells in an epiblast-like state, with reduced differentiation towards endoderm and mesoderm prior to the aggregation stage. Since cell density increases over the culture period, we performed pre-CHIR treatment one-day earlier, at day 3 of culture to examine gene expression in both A3FX (Fig. S2A) and A20FX (Fig. S2B). Notably, TBXT and FOXA2 expressing cells were distributed across colonies at all cell densities in A3FX, suggesting that TBXT and FOXA2 induction depend on cell density upon CHIR addition (Fig. S2A). On the other hand, FOXA2 was not strongly induced in A20FX culture at day 3 pre-CHIR treatment (Fig. S2B). Since cell density determines the cell fate choice between surface ectoderm and amnion from hPSC (Nakanoh, 2025; Nakanoh et al., 2024), our findings provide further insight to develop efficient differentiation strategies to expand specific lineages from hPSCs. Taken together, these results demonstrate that the generation of elongated gastruloids via pre-CHIR treatment is critically dependent on using vigorous hPSC colonies at an optimal cell density.

### Gastruloid generation without pre-CHIR treatment

So far, our data suggest that hPSC cultures post-CHIR treatment are heterogeneous for endoderm, mesoderm and neuronal stem cell states, and that these states are dynamic and sensitive to cell density and other conditions. We considered that aggregation from these mixed cell populations may contribute to stochastic variability of gastruloids, and that CHIR itself may be contributing to this heterogeneity. To address this, we attempted to generate gastruloids without pre-CHIR treatment (Fig. 3A), inspired by other related organoid protocols (Meijer et al., 2025; Rito et al., 2025; Sanaki-Matsumiya et al., 2022). Primed (p)HNES1 cells were cultured in both A3FX and A20FX, resulting in no apparent difference in the morphology (Fig. 3B), but a slight increase in FOXA2+ cells was observed in A3FX cultures (Fig. 3C). Using A3FX and direct aggregation, elongating gastruloids could be formed more efficiently than those cultured in A20FX (Fig. 3D). A3FX-derived gastruloids elongated consistently compared with those grown in A20FX and expressed the expected lineage markers: TBXT and SOX2 at the posterior end with GATA6 at the anterior (Fig. 3E), an expression pattern similar to that of pre-CHIR treated gastruloids (Fig.1). Notably, gastruloids differentiated by direct aggregation exhibited reduced variation and higher efficiency than those cultured in A20FX (Fig. 3F).

**Figure 3.**
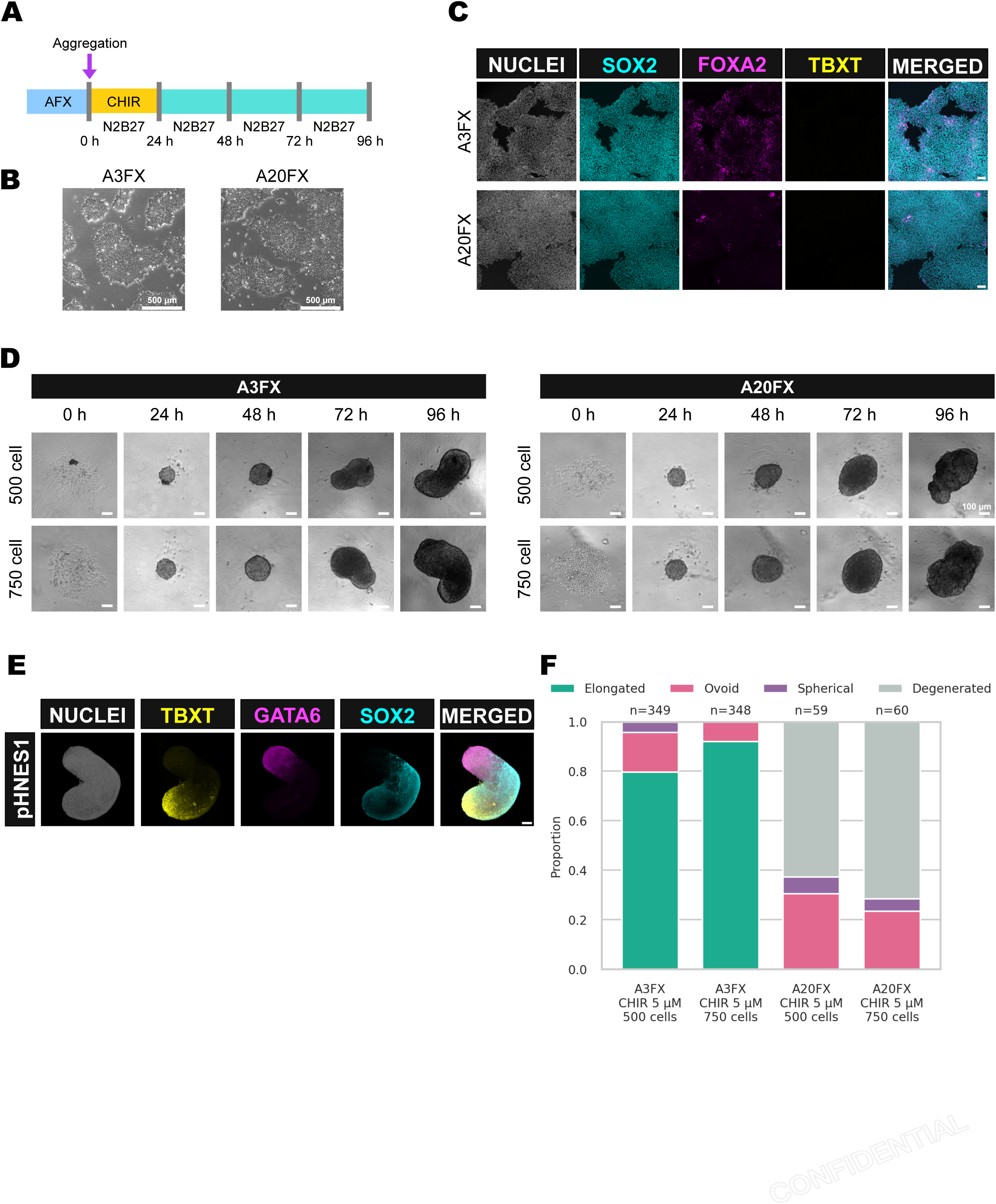
Gastruloid differentiation via direct aggregation protocol. (A) Schematic for the protocol for making gastruloids without pre-CHIR treatment. (B) Representative bright field images for A3FX and A20FX culture pHNES1 hPSCs before making gastruloids. Scale bar = 500 µm. (C) Immunofluorescence for SOX2, FOXA2, and TBXT in A3FX and A20FX culture pHNES1 hPSCs. Scale bar = 100 µm. (D) Representative images of gastruloid differentiation over the time course. Scale bar = 100 µm. (E) Immunofluorescence for SOX2, GATA6, and TBXT of gastruloids generated from A3FX cultured pHNES1 hPSCs. Scale bar = 100 µm. (F) Proportion of elongated, ovoid, spherical, and degenerated morphology from pHNES1 hPSCs. Plot represents a pool of 12 (A3FX) and 3 (A20FX) independent experiments. The number of gastruloids (n) for each condition is indicated above the corresponding plot.

To determine whether WNT activation is perturbated by cell density, we stained cultures for the WNT target, LEF1, and observed strong induction at low density in A3FX (Fig. S3A), but expression was suppressed in high density A20FX cultures (Fig. S3B). Interestingly, nuclear localized active *β*-Catenin accumulated in A20FX culture, but was suppressed in high density after pre-CHIR treatment, suggesting the balance between TBXT+, FOXA2+, and SOX2+ are determined by WNT activity (Fig. S4).

To improve the efficiency of gastruloid elongation, we added TGF*β*-inhibitor SB431542 (SB43) after aggregation, since SB43 was previously employed to generate gastruloids from H9 hPSCs (Hamazaki et al., 2024) (Fig. 4A). SB43 significantly improved the efficiency of elongated gastruloid formation from H9 hPSCs cultured in A3FX. In A20FX, most aggregates became ovoid when SB43 was added (Fig. 4B and C). Interestingly, although the consistently of shape and integrity of the H9 gastruloids was improved with SB43, GATA6 expression was absent (Fig. 4D), suggesting that transient TGF-*β* inhibition may promote posterior patterning in gastruloids, a possibility that warrants further investigation.

**Figure 4.**
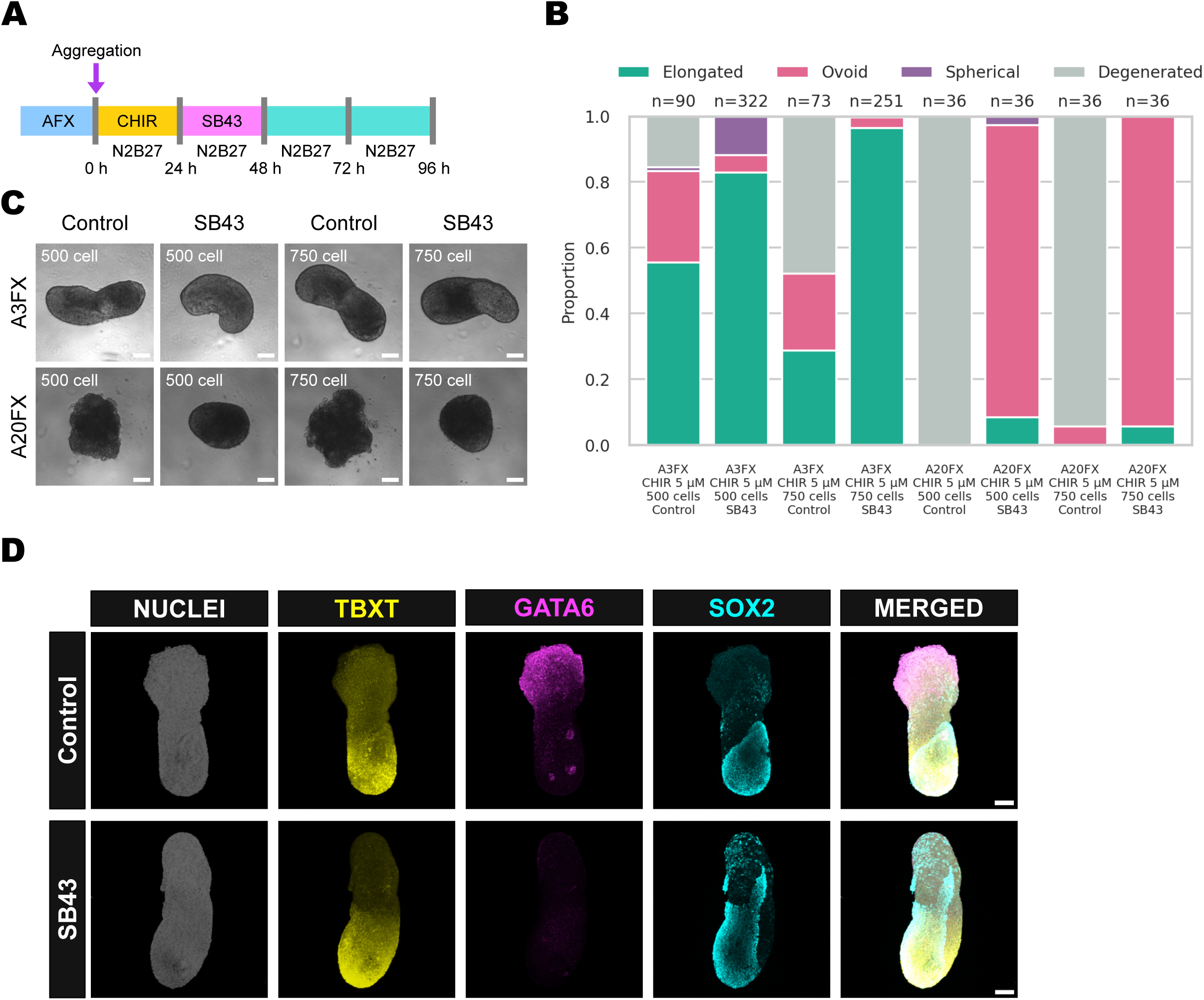
TGF*β* inhibition improve gastruloid differentiation in H9. (A) Schematic for the protocol for making gastruloids without pre-CHIR treatment and timed TGF*β* inhibition. TGF*β* inhibitor SB43 was added between 24h and 48h. (B) Proportion of elongated, ovoid, spherical, and degenerated morphology from pH9 hPSCs generated from the indicated condition. The number of gastruloids (n) for each condition is indicated above the corresponding plot. The plots represent data pooled from 5 (A3FX control), 10 (A3FX SB43), 3 (A20FX control), and 3 (A20FX SB43) independent experiments. (C) Representative images of gastruloid differentiation over the time course. Scale bar = 100 µm. (D) Immunofluorescence for SOX2, GATA6, and TBXT of gastruloids generated with or without AB43 from A3FX cultured pH9 hPSCs. Scale bar = 100 µm.

In conclusion, by defining the key parameters governing human gastruloid formation, we have developed a simplified and robust protocol for efficient and consistent generation of elongated gastruloids from human PSCs (Fig. S5). This streamlined method improves accessibility and reproducibility, providing a more controlled platform to investigate fundamental principles of early human development.

## Materials and methods

### Ethics statement

The human cell lines used in this study comprise H9, one of the first PSC lines to be derived directly from blastocysts in USA (Thomson et al., 1998), which has been widely used by the stem cell and developmental biology community, and HNES1 (Guo et al., 2016), derived in UK from a disaggregated isolated ICM from a blastocyst six days old, donated to Human Fertility and Embryology Authority (HFEA) licence R0178 with informed consent from patients undergoing assisted conception procedures, for purposes, including derivation of pluripotent stem cell lines stipulated in the licence application and patient information document. The gastruloids generated from these cells lack extra-embryonic tissues, anterior structures and circulating blood; they therefore pose little risk of acquiring organised brain tissue or sentience and consequently require no specific ethical approval.

### hPSC culture

H9 and primed HNES1 hPSCs were used in this study. Naïve HNES1 were capacitated to the primed state according to the previously reported protocol (Rostovskaya et al., 2019). All hPSCs were cultured in N2B27 supplemented with 3 ng/ml or 20 ng/ml Activin A (Qkine), 10 ng/ml FGF2 (Qkine), and 2 µM XAV939 (Tocris) on 1:100 diluted Geltrex (Gibco) coated plates at 37°C, 7% CO_2_, 5% O_2_. hPSCs were passaged every 4 days using Accutase (BioLegend). ROCK inhibitor Y-27632 (Tocris) was used at 10 µM when passaging hPSCs.

### Conventional human gastruloid induction

Conventional human gastruloids were induced according to previous reports with modifications (Moris et al., 2020). 2x, 4x, 6x, and 8x10^4^ hPSCs were seeded on 1:100 diluted Geltrex coated 12-well plate in N2B27 with A3FX or A20FX and 10 µM Y-27632. Between day1 and day3, the medium was changed to N2B27 with AF3X or A20FX. On day 4, the medium was replaced to N2B27 supplemented with 3 ng/ml or 20 ng/ml Activin A, 10 ng/ml FGF2, and 4 µM CHIR99021. After 24h, cells were dissociated into single cells with Accutase. Then, 500 cells were plated into round bottom 96-well plate with 40 µL N2B27 with 4 µM CHIR99021, CEPT cocktail (Chen et al., 2021; Ryu et al., 2023; Tristan et al., 2023). After 24h, 150 µl N2B27 was added to each well. From 48h, 150 µl of medium was removed and 150 µl N2B27 was added to each well by multi-channel pipette. Gastruloids were cultured until 96h.

### Gastruloid induction with direct aggregation

hPSCs were cultured with N2B27 with A3FX on 1:100 diluted Geltrex coated plate for 4 days until forming compacted colony with about 80% confluency. Cells were dissociated into single cells with Accutase. Then, 500 or 750 cells were plated into round bottom 96-well plate with 40 µL N2B27 with 4-5 µM CHIR99021 and CEPT cocktail. After 24h, 150 µl N2B27 with or without 10 µM SB431542 (Tocris) was added to each well. From 48h, 150 µl of medium was removed from each well and 150 µl N2B27 was added to each well by multi-channel pipette. Gastruloids were cultured until 96h.

### Immunofluorescence

For immunofluorescence of hPSCs in 2D, cells were cultured on plastic cover slips coated with 1:100 diluted Geltrex. Cells were fixed at the indicated time points in 4% PFA for 15 min at room temperature (RT), followed by washing in PBS. Cells were permeabilized in 0.1% TritonX-100/PBS for 10 min at RT. Blocking reaction was carried out with 5% donkey serum, 5% BSA, and 0.1% TritonX-100 in PBS for 1 hour at RT. Primary antibodies were diluted in blocking solution and incubated overnight at 4℃. After washing cells in 0.1% Tween 20 in PBS, cells were incubated with Alexa Fluor-conjugated secondary antibodies (Thermo) for 1 hour at RT. Nuclear staining was carried out with Hoechst33342 (Thermo).

Gastruloids were collected from each stage and fixed in 4% PFA for overnight at 4℃, followed by washing in PBS. Gastruloids were permeabilized in 0.5% Triton X-100 in PBS for 15 min and blocked with 5% donkey serum, 5% BSA, and 0.1% Tween20 in PBS for overnight at 4℃. Primary antibodies (Table S1) were diluted in blocking solution and incubated overnight at 4℃. After washing gastruloids in 0.1% Tween 20 in PBS, gastruloids were incubated with Alexa Fluor-conjugated secondary antibodies (Thermo) for 2 h at RT. Nuclear staining was carried with Hoechst33342 (Thermo).

### Imaging and image analysis for 2D culture cells

Bright field images for gastruloids were acquired using EVOS M7000 imaging system (Invitrogen). Fluorescent images for gastruloids were acquired using Stellaris 8 confocal microscope (Leica) and processed with ImageJ/Fiji. Nuclear segmentation was performed using Cellpose3 (Stringer et al., 2021). The resulting regions of interest (ROIs) were transferred to each channel in Fiji, and the mean intensity of each cell was quantified using the “Measure” function. The expression level of each gene was normalized to the corresponding Hoechst channel intensity and plotted. To compare expression levels between the colony center and periphery, intensity profiles across colonies were quantified using the “Plot Profile” function in Fiji. The resulting intensity for each channel was then visualized as a line plot.

### Imaging and image analysis for gastruloids

Bright field images for gastruloids were acquired using EVOS M7000 imaging system (Invitrogen). Fluorescent images for gastruloids were acquired using Stellaris 8 confocal microscope (Leica) and processed with Fiji.

## Acknowledgements and funding

We are indebted to the pioneers of gastruloid technology, particularly those rising to the challenge of developing protocols for use with human pluripotent stem cells. We wish to thank our colleagues, Lawrence Bates, Kasia Milto, Ewa Ozga and Anastasiia Bekhtereva for discussion and utilisation of gastruloid technology. We are grateful to Ann Wheeler and her team for their valuable contribution to advanced microscopy and image analysis. This work was funded by Wellcome Trust Collaborative Award 220379/B/20/Z and the University of Edinburgh.

---

**Supplementary Figure 1.**
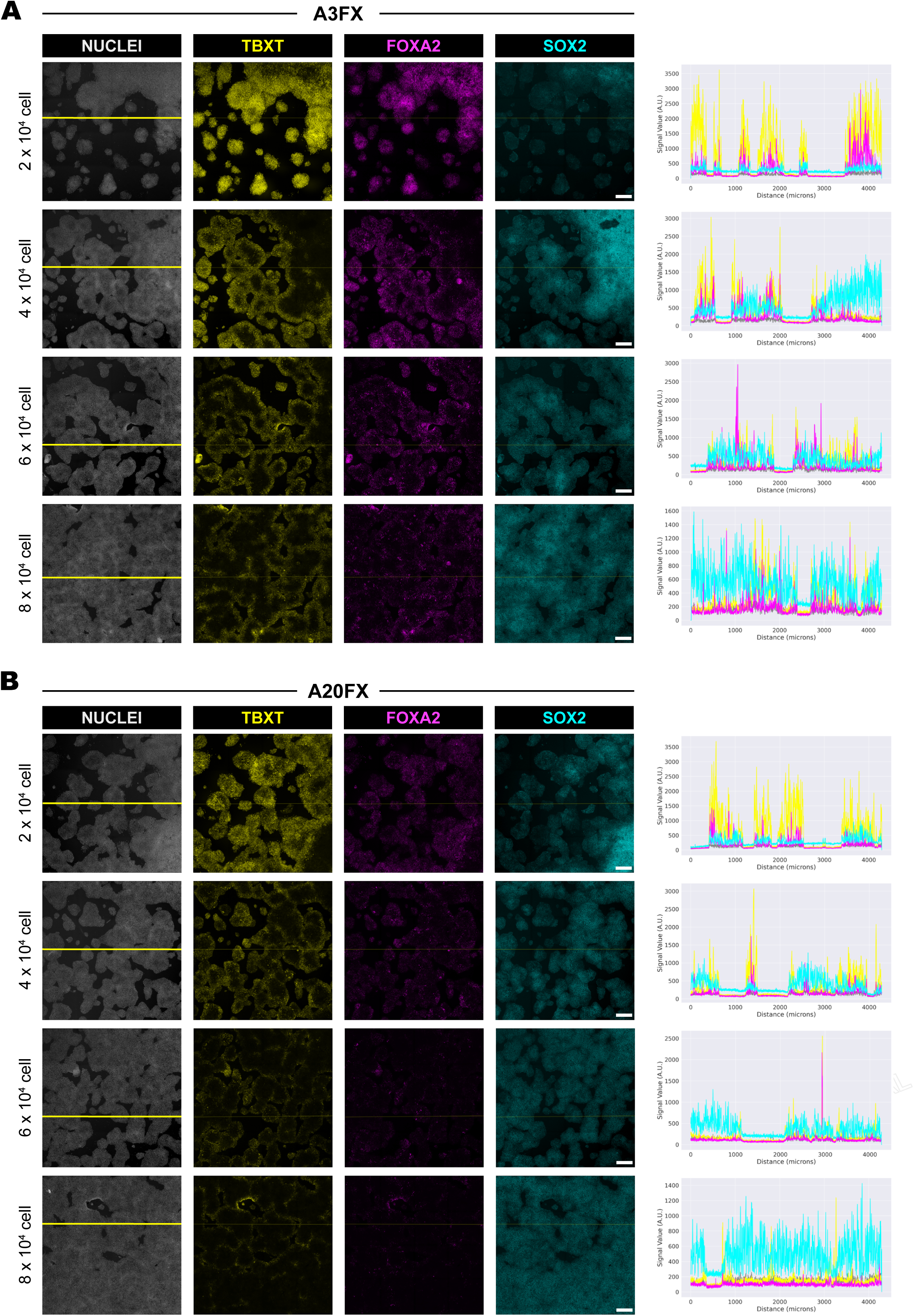
(A, B) Immunofluorescence for TBXT, FOXA2 and SOX2 in pre-CHIR treated pHNES1 hPSCs from A3FX (A) or A20FX (B). pHNES1 hPSCs were plated at the indicated cell number in 12-well plate and cultured for 4 days in A3FX or A20FX; pre-CHIR treatment was performed with 4 µM CHIR/A3F or A20F for 24 h. The image was generated by stitching together a 5x5 tile scan. Scale bar = 500 µm. To compare expression at the colony centre and periphery, signal intensities for TBXT, FOXA2, and SOX2 were measured along the yellow line (nuclei channel) using Plot Profile and visualized as a line plot.

**Supplementary Figure 2.**
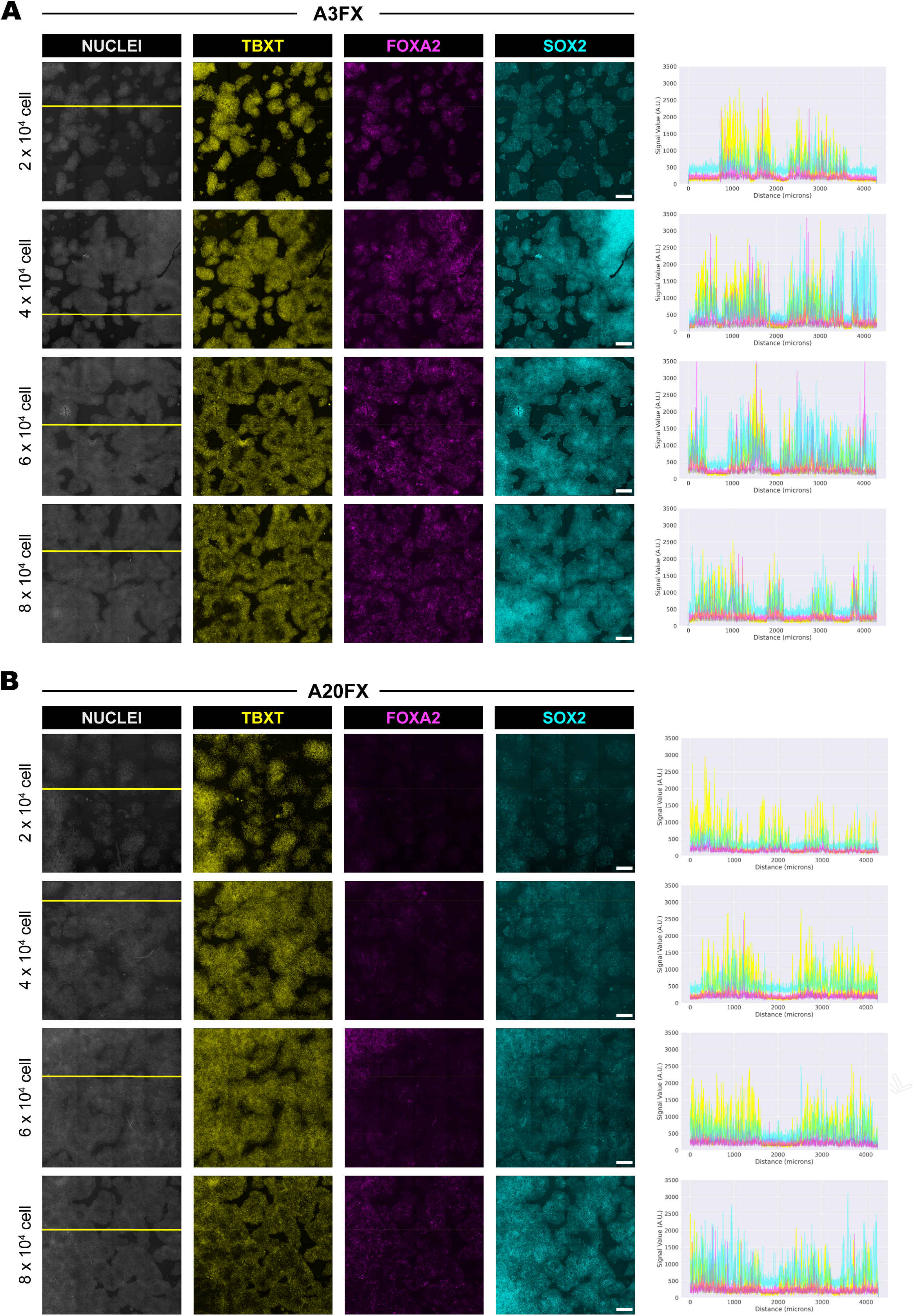
(A, B) Immunofluorescence for TBXT, FOXA2, and SOX2 in pre-CHIR treated pHNES1 PSCs from A3FX (A) or A20FX (B). pHNES1 hPSCs were plated at the indicated cell number in 12-well plate and cultured for 3 days in A3FX or A20FX, and pre-CHIR treatment was performed with 4 µM CHIR/A3F or A20F for 24 h. The image was generated by stitching together a 5x5 tile scan. Scale bar = 500 µm. To compare expression at the colony centre and periphery, signal intensities for TBXT, FOXA2, and SOX2 were measured along the yellow line (nuclei channel) using Plot Profile and visualized as a line plot.

**Supplementary Figure 3.**
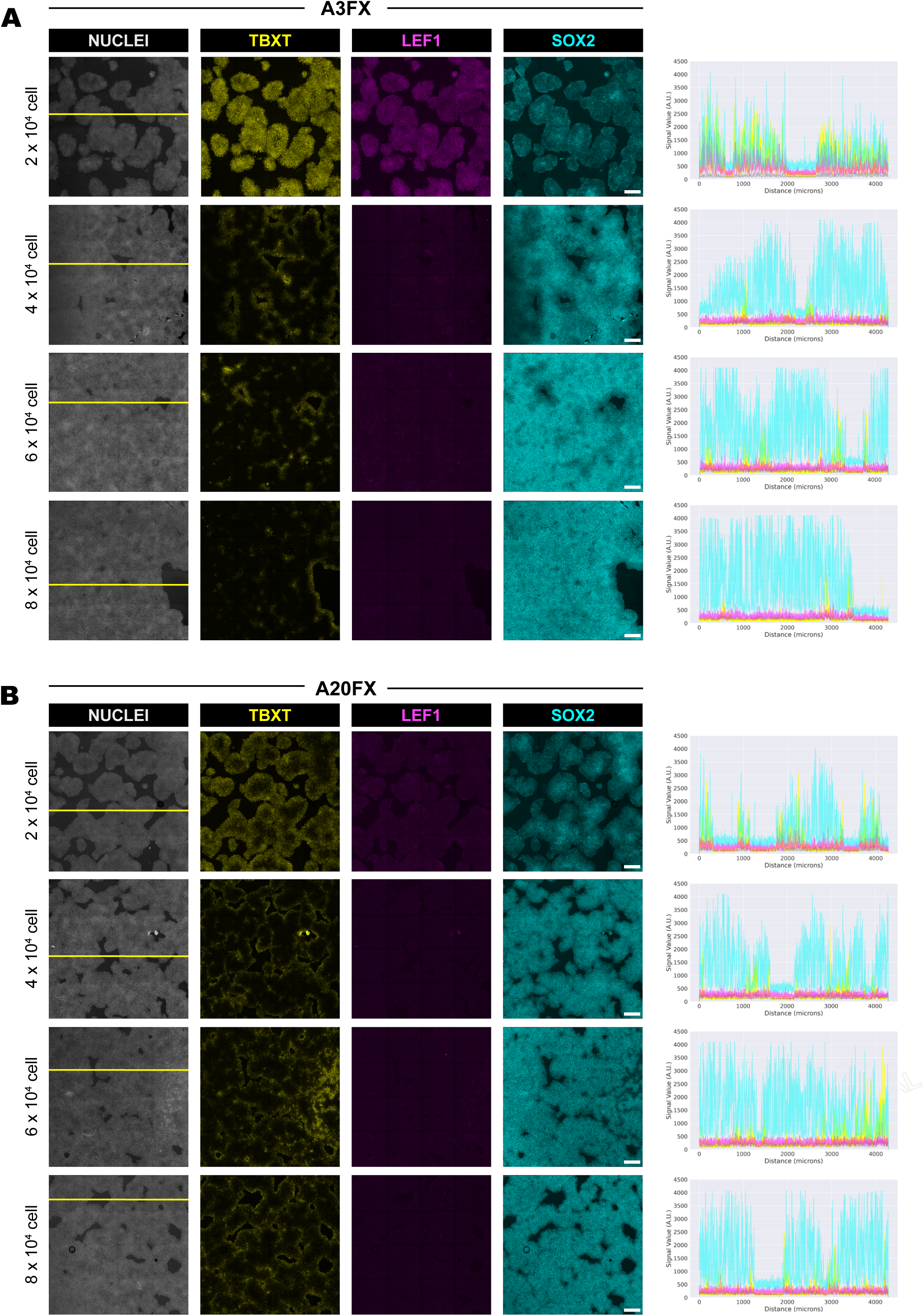
(A, B) Immunofluorescence for TBXT, LEF1, and SOX2 in pre-CHIR treated pHNES1 hPSCs from A3FX (A) or A20FX (B). pHNES1 hPSCs were plated at the indicated cell number in 12-well plate and cultured for 4 days in A3FX or A20FX, and pre-CHIR treatment was performed in 4 µM CHIR/A3F or A20F for 24 h. The image was generated by stitching together a 5x5 tile scan. Scale bar = 500 µm. To compare expression at the colony centre and periphery, signal intensities for TBXT, LEF1, and SOX2 were measured along the yellow line (nuclei channel) using Plot Profile and visualized as a line plot.

**Supplementary Figure 4.**
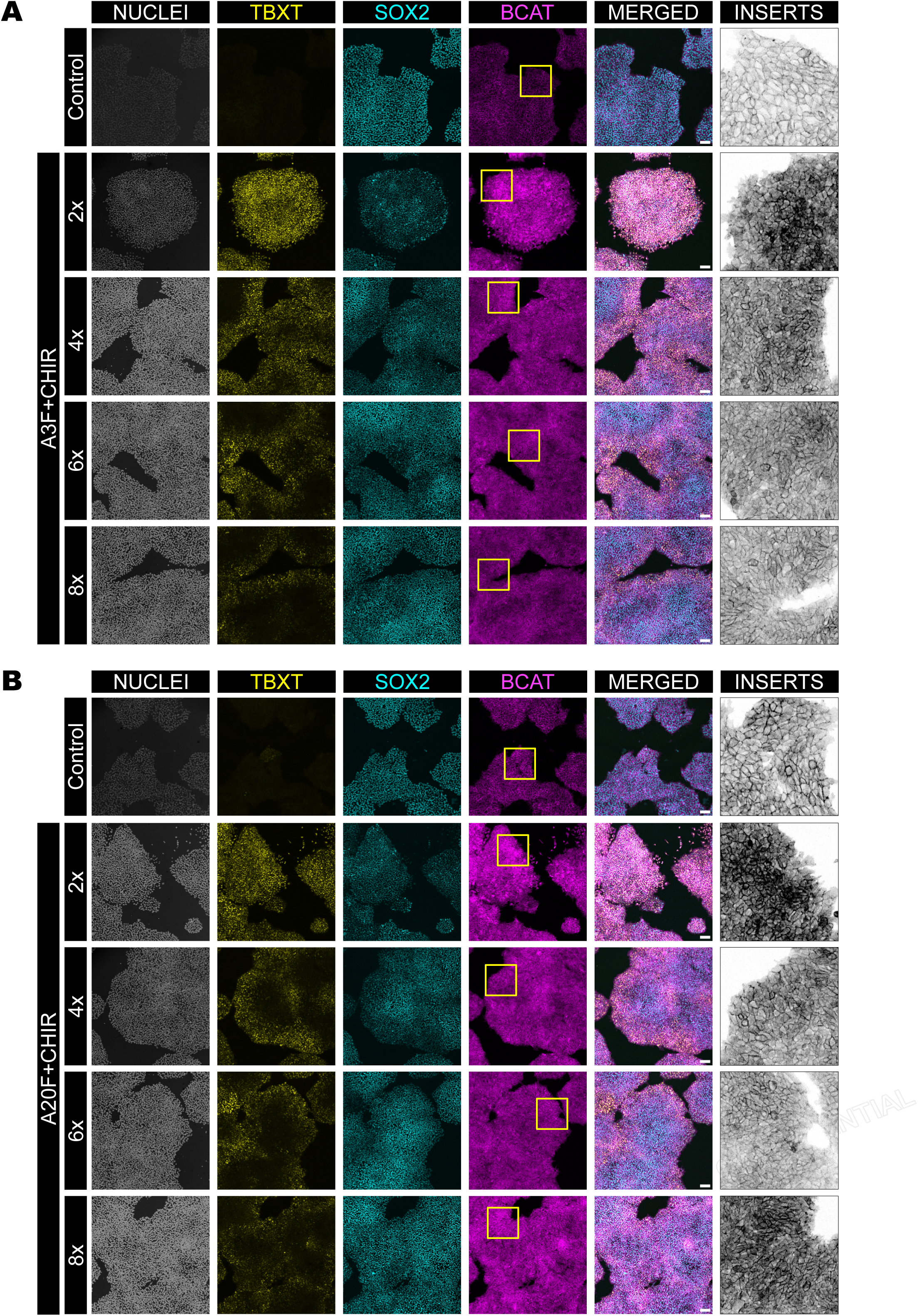
(A, B) Immunofluorescence for TBXT, SOX2, and *β*-Catenin (BCAT) after pre-CHIR treated pHNES1 hPSCs in A3FX (A) or A20FX (B). pHNES1 hPSCs were plated at the indicated cell number in 12-well plate and cultured for 4 days in A3FX or A20FX, and pre-CHIR treatment was performed in 4 µM CHIR/A3F or A20F for 24 h. Scale bar = 100 µm. The inset shows a magnified view of the region outlined by the yellow box in the corresponding *β*-Catenin-stained image for each condition.

**Supplementary Figure 5.**
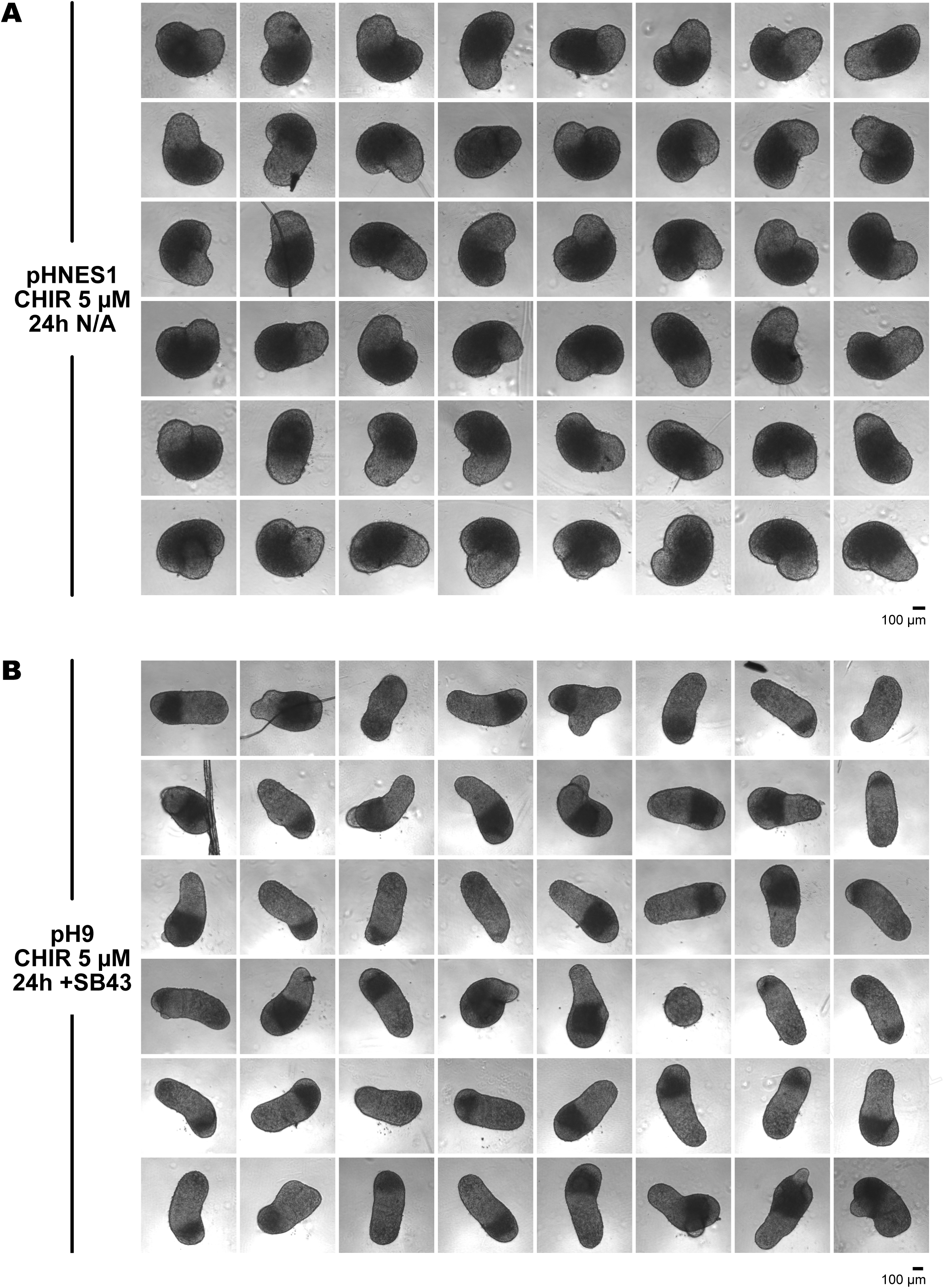
(A, B) Bright field images of gastruloids generated from pHNES1 (A) and pH9 (B) hPSCs with indicated condition using direct aggregation protocol. Scale bar = 100 µm.

**Supplementary Table 1.** List of antibodies used in this study.

| **Primary antibodies** | **Source** | **Cat#** |
| --- | --- | --- |
| Rat anti-SOX2 | eBioscience | 14-9811-82 |
| Rabbit anti-FOXA2 | abcam | ab108422 |
| Goat anti-Brachyury (TBXT) | R&D | AF2085 |
| Rabbit anti-GATA6 | Cell Signaling | 5851 |
| Rabbit anti-LEF1 | Cell Signaling | 2230 |
| Rabbit anti-beta Catenin | abcam | ab16051 |

